# Architecting a distributed bioinformatics platform with iRODS and iPlant Agave API

**DOI:** 10.1101/034488

**Authors:** Liya Wang, Peter Van Buren, Doreen Ware

## Abstract

Over the past few years, cloud-based platforms have been proposed to address storage, management, and computation of large-scale data, especially in the field of genomics. However, for collaboration efforts involving multiple institutes, data transfer and management, interoperability and standardization among different platforms have imposed new challenges. This paper proposes a distributed bioinformatics platform that can leverage local clusters with remote computational clusters for genomic analysis using the unified bioinformatics workflow. The platform is built with a data server configured with iRODS, a computation cluster authenticated with iPlant Agave system, and web server to interact with the platform. A Genome-Wide Association Study workflow is integrated to validate the feasibility of the proposed approach.

## I. Introduction

Massive amount of data produced with the rapidly advancing next generation sequencing technologies [1] has become a major challenge to traditional computational infrastructures. Cloud computing (“pay-per-use model”) is believed to be a scalable and cost efficient solution to the big data challenge and many platforms and workflow systems have been proposed [2, 3]. On the other hand, many institutes have their own genomic sequencing center and analysis clusters for handling large amount of data analysis locally. When more than one institutes or sequencing centers are working on one large consortium project, efficient data coordination becomes a new challenge. Completely replying on cloud computing will waste local computing resources and in turn increase costs on both computation and storage.

Therefore, a hybrid system that leverages local and remote resources, allows automatic data and metadata synchronization, and supports unified bioinformatics workflow, will be ideal for large-scale cross-institute collaborations. To address these problems, we propose a distributed bioinformatics platform built at Cold Spring Harbor Laboratory (CSHL) for optimizing data transfer, management, and analysis among local and remote systems. The platform is built with one computing cluster authenticated with iPlant Agave (or “A grid and virtualization environment”) system [4], one data Resource server configured with iRODS (or “integrated Rule-Oriented Data System”) [5] zone of the iPlant Collaborative project [6].

The remainder of this paper is structured as follows: Section II briefly introduces iRODS; Section III briefly introduces the iPlant project and particularly the Agave API; Section IV provides an overview of the system and dataflow through the platform for bioinformatics analysis; Section V presents integration of a dozen Genome Wide Association Study (GWAS) applications (or tools, apps) into the system; Finally, conclusion and future directions are presented in Section VI.

## II. iRODS

iRODS is open source data management software used by research organizations and government agencies worldwide. It uses a metadata catalog for easy data discovery and allows access to distributed storage assets through data virtualization. It automates data workflows with a rule engine that permits any action to be initiated by any trigger on any server or client in the grid and enables secure collaboration for easily accessing data hosted on a remote grid.

There are two types of iRODS server in an iRODS Zone (an independent iRODS system), iCAT and Resource: The iCAT server handles the database connection to the iCAT metadata catalog. It can also provide storage resource. One iRODS Zone will have exactly one iCAT server; The Resource server provides additional storage resources. One iRODS Zone can have zero or more Resource servers.

## III. iPlant

The iPlant Collaborative project is funded by National Science Foundation to create an innovative, comprehensive, and foundational cyber-infrastructure (CI) in support of plant biology research. iPlant is an open-source project that allows the community to extend their infrastructure to meet different needs.

### A. Data Store

The iPlant Data Store is a federated network of iRODS servers running at University of Arizona and mirrored at the Texas Advanced Computing Center (TACC). iPlant users can access the Data Store in multiple ways, including iPlant Discovery Environment (DE, a web-based data and analysis platform) [7], iDrop [8], Cyberduck [9], RESTful web service interface through iPlant Agave API, iCommands from iRODS, and FUSE interface [10] for command line tools.

### B. Agave API

Agave is an open source, platform-as-a-service solution for hybrid cloud computing. It is developed by the iPlant project to provide a full suite of services covering everything from standards-based authentication and authorization to data computational, and collaborative services.

Agave provides an existing catalog with hundreds of today’s most popular scientific apps. To deploy a new app that runs on the command line, e. g. source code, binary code, VM images, Docker images [11], or mix and match with apps from the catalog, user simply needs to host them on a storage system registered with Agave (e. g. the iPlant Data Store), and provide Agave a JSON (JavaScript Object Notation) file which describes path to the executable files, the execution system, and tool information (inputs, outputs, and parameters). The app can then be executed either through Agave directly (on the command line), or through DE (with an automatically generated graphical user interface (GUI) for each app).

User can register two types of system with Agave: storage and execution system. Agave supports authentication through password, ssh keys, or ssh tunnels, data transfer through ftp, sftp, gridftp, iRODS, or Amazon S3, and scheduler including SGE, SLURM [12], or Condor [13], for the execution system.

## IV. Overview of the Distributed System

Fig. 1 shows the architecture of our distributed system and how the system interacts with remote systems at University of Arizona (UA) and TACC. There are two important nodes: 1) The CSHL storage server provides storage space for storing massive amount of data locally. The server is enrolled into iPlant’s iRODS system as a Resource server, which makes the data manageable and accessible to scientific workflows inside iPlant DE (top left icon). Resource server hold not only input data but also executable files of all scientific workflows. The resource server at CSHL is registered as a storage system to the Agave system residing at TACC through Agave API. This allows the Agave service to bring the input data and executable files to an execution system at runtime. Additionally, data replications among resource servers at UA, TACC, and CSHL are enabled through iRODS; 2) The CSHL cluster node provides computation resource for local data. It is registered as an execution system with Agave system through Agave API. In addition, we set up a web server [14] to enable direct communication with user, storage and computer servers. The web server is also registered with Agave as a storage system for storing analysis results.

**Figure 1.**
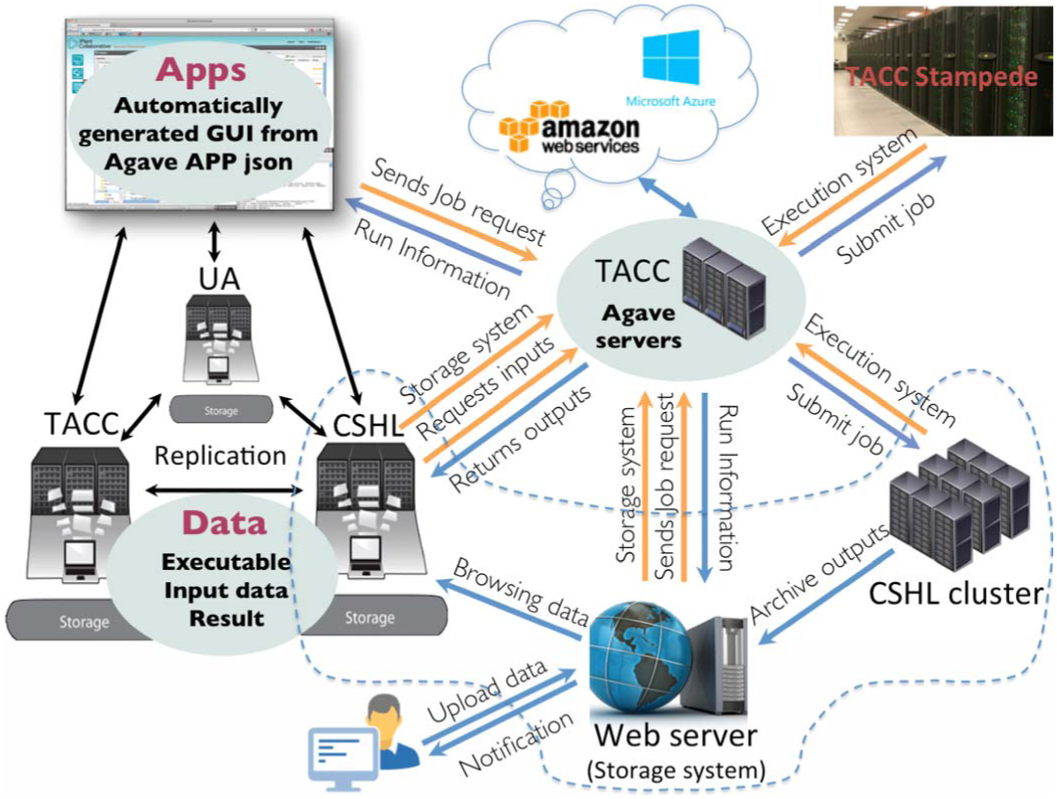
Architecture of the CSHL system (enclosed with dashed line) and interactions with systems at UA and TACC

In summary, inputs, analysis results, and executable files can be hosted or transferred after completion of an analysis to the web server, storage server, execution server, or even cloud resources such as Amazon S3/EC2 or Microsoft Azure if they are registered with the Agave system. A workflow can be easily pointed to any systems for retrieving/storing data or performing large-scale computation.

## A. Resource Server

The CSHL storage server (Thinkmate STX-EN XE24-2460 with 100 TB raw storage) is added as an additional Resource server to the iPlant Zone with following steps:

- Create user account for irods: useradd irods
- Download irods an unpack the gzipped file: https://wiki.irods.org/index.php/Downloads
- Go to the directory iRODS and run./irodssetup
- Enter iplant as iRODS zone name and pick a name for the resource server during setup:

Besides adding the storage server as a Resource server, another option is to configure the storage server as an iCAT server then follows the federation manual [15] to communicate with another iRODS zone (e.g. iPlant zone in this example). we choose to add it as a Resource server to simplify the daily management of a standalone iRODS system. However, for projects involving more sophisticated data operation, federation will be more appropriate.

Like other institutes, servers at CSHL are protected from internet traffic by a firewall appliance. In order to allow iRODS servers to communicate, firewall exception rules were created on standard iRODS tcp ports to specific IP addresses: tcp/1247 and tcp/20000-20399. Specifically, files smaller than 32 MB are transferred through iRODS control port 1247; For large files, iRODS catalog server directs a server to open parallel ports 20000-20399 that enables fast data transfer.

Once setup is complete, we benchmarked the data transfer rate between CSHL cluster and CSHL resource server and compared them with the rate between CSHL cluster and UA resource servers. The testing is done with iCommands (iget or iput with –R specifying the specific resource server) and results are listed in Table I, which shows that it can be more than 8 times faster to bring the data to cluster for analysis if the data is stored locally.

**TABLE 1.**
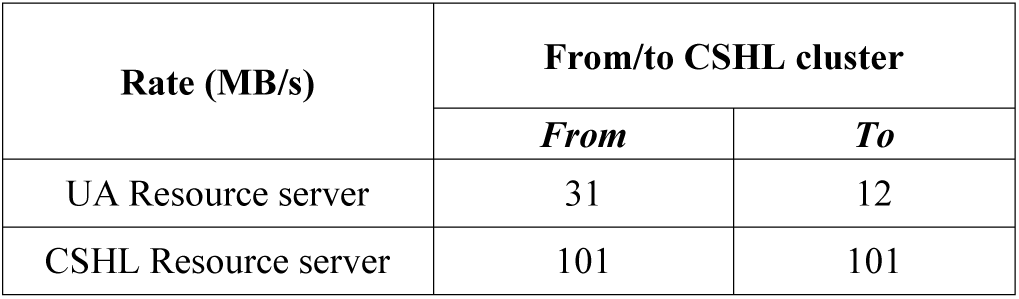
Data transfer rate

| Rate (MB/s) | From/to CSHL cluster |  |
| --- | --- | --- |
|  | From | To |
| UA Resource server | 31 | 12 |
| CSHL Resource server | 101 | 101 |

### B. Computing Cluster

The computing cluster is a Dell PowerEdge R820 server. It is a four sockets server with 48 physical CPU cores, 1 TB of memory and 10 TB of local storage. CentOS 6 is installed. A standard Linux installation was performed and a LVM logical volume was created for a data partition. SLURM is chosen for its simpler daemons architecture than Condor given that SLURM is designed to handle mostly homogeneous clusters.

Before authenticating with Agave, role user account is created on the server, which is the acting account that submits the analysis jobs to the cluster. Then we store the cluster information in a JSON file and register it with Agave via following command:

> $ systems-addupdate –F cshl-compute-01.json

The system JSON file contains the system id, how Agave worker access the cluster, scheduler, queue, resource limitation, etc. For resource management, we also use sacctmgr from SLURM to enforce fairshare priority.

The Agave execution system can be shared with selected iPlant users or made public to all users. By itself, this will enable sharing the computing clusters across institutes if needed. In addition, we installed Docker image of Agave-client [16] so we can interact with the Agave system directly from the CSHL cluster.

### C. Dataflow

To better illustrate how the system functions for a typical bioinformatics analysis, Fig. 2 describes a summary of the dataflow, with more details explained below.

**Figure 2.**
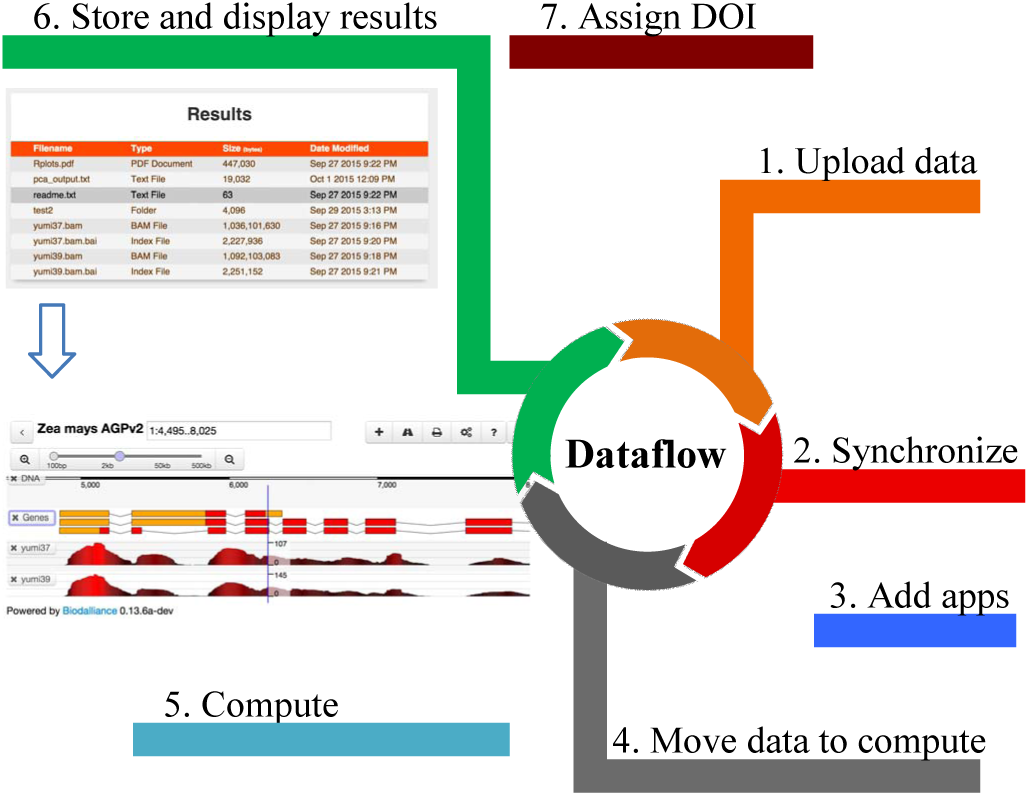
Data flow for typical bioinformatics analysis using the distributed system

#### 1) Upload data

In the data window of iPlant DE, user can import files from URLs or upload multiple small files (<1.9 GB each) from their desktops directly. For large files, users are directed to using iCommands (command line) or Cyberduck (with GUI). For both iCommands and Cyberduck, user can specify the resource server where the data is expected to go.

The CSHL web server [14] supports both direct uploading and importing from URL. Currently, the uploaded data is hosted right on the web sever and brought to the compute server when an analysis job is submitted. For large-scale data sets, we use iCommands to upload them to the CSHL resource server and they are available in iPlant DE immediately once uploaded.

#### 2) Synchronize data

As mentioned earlier, data are synchronized through iRODS replication. The replication can be setup to execute when network traffic is low, e.g. weekend or late night. As of November 2015, about 1.26 PB data have been synchronized between UA and TACC. The automated synchronization improves data safety with extra copies, and more importantly, it gives users the flexibility to analyze their data on different clusters (UA, TACC, or CSHL) with unified workflows.

Even if users choose to not replicate their data to different resource servers, the data is still manageable inside iPlant DE. Therefore, within iPlant DE, user can add metadata/comments or apply various metadata templates to the raw data. The metadata is searchable and can also be dumped out and consumed by third party systems.

#### 3) Add analysis tool

More details on how to integrate an Agave app can be found in section V. For Agave apps, to ensure consistency among different execution systems, all executable files are hosted in a registered storage system (e.g. iPlant Data Store) and copied to the execution system at runtime. As a side note, iPlant DE also supports tool integration through Docker besides Agave.

#### 4) Move data to execution system

Data are moved to the execution system at runtime when an analysis job is submitted. Depending on the requirements of CPU and memory, status of queues, where the data is physically located, user can choose the local or remote execution systems by re-pointing the Agave apps to them. Re-pointing can be easily done by following steps: a) retrieve a particular app’s JSON definition from the Agave system; b) modify the execution system definition in the JSON file; and c) register the new JSON file as a new app to the Agave system.

#### 5) Compute

An analysis can be parallelized differently depending on the execution system it is running on. For embarrassingly parallel problem, the parametric launcher module on TACC clusters or SLURM job arrays on the CSHL clusters can be executed through a wrapper script shipped with the app.

#### 6) Store and display results

For the CSHL system, we choose to register the web server as a storage system so that we can use Agave to archive analysis results back to the web server. A landing page is automatically generated for viewing the results folder. The landing page also attaches link to specific resulted file for visualizing them directly on a Genome Browser (Fig. 2), or other JavaScript based web visualization tools (e.g. Manhattan plot for GWAS etc.). The URLs for the results can also be used as inputs directly for subsequent analysis.

#### 7) Assign DOI

The results page can be shared with collaborators directly. For permanent identifiers, iPlant assigns DOIs for data, workflow, project, and even a customized virtual machine if required metadata are provided.

## V. Integrating apps

To validate the feasibility of the proposed approach, we converted a dozen iPlant apps [17] and made them publically available through both iPlant DE and CSHL web server [14]. The conversion is done by modifying the JSON file describing each app and re-publishing them with Agave by following command:

> $apps-addupdate –F mytool.json

The modification includes updating the execution system (from TACC cluster to CSHL cluster) and change queue names (if different). So all iPlant Agave apps can be easily converted to run on CSHL cluster with the exception of those relying on TACC modules (pre-built software packages). One solution we adopted for these apps is to install those packages on the CSHL server when needed.

Instructions for deploying a brand new app via Agave are available on Github [18]. Briefly, besides the JSON file describing the app, a shell script is needed for each app to convert the parameters and inputs from the JSON file for command line execution.

## VI. Conclusions and Future Work

In this work, we describe the architecture of a distributed system to leverage local servers with remote resources. By making a dozen CSHL system powered GWAS apps available both in iPlant DE and on CSHL web server, we demonstrate that the system allows easily sharing of data, apps, and workflows across institutes. This will be critical for collaborations involving multiple sequencing centers. With such system, data can be processed locally and shared among each other, which will reduce data traffic across the nation, enable easy data management and sharing, and reduce the cost of cloud computing if needed.

As shown in Fig. 1, we have built a web server to interact with user, storage and computing server directly, a prototype of the web service is available at [14], which exposes the GWAS apps integrated in this paper as an alternative web service in addition to iPlant DE. For future developments, we are focusing on automating metadata integration from Agave and iRODS system, further optimizing local data transfer and management, developing more visualization tools, and providing web services to third party biological databases powered by both local and cloud resources. More importantly, we will be coordinating with the iPlant team closely to leverage iPlant Data Common efforts on developing a more sophisticated metadata management system.

## Acknowledgment

This work is supported by the National Science Foundation Plant cyber-infrastructure Program (#DBI-0735191 and DBI-1265383). The authors would also like to thank Drs. Rion Dooley and Matthew Vaughn on integrating the CSHL cluster with the Agave system, Dr. Nirav Merchant, Andy Edmonds, and Edwin Skidmore on integrating the resource server with iPlant iRODS zone, Zhenyuan Lu, Kapeel Chougule, and Cornel Ghiban for setting up the prototyped web server.

